# Sperm and northern bottlenose whale interactions with deep-water trawlers in the western North Atlantic

**DOI:** 10.1101/2021.10.25.464263

**Authors:** Usua Oyarbide, Laura Feyrer, Jonathan Gordon

**Author notes:** **Corresponding author:** (UO).

## Abstract

Commercial fisheries have increased in all the world’s oceans with diverse unintended impacts on marine ecosystems. As a result of resource overlap, interactions between cetaceans and fisheries are a common occurrence and, in many cases, these give rise to significant conservation issues. Research on the distribution and types of such interactions is important for efficient management. In this study, we describe the behaviors of two whale species: sperm whales (*Physeter macrocephalus*) and northern bottlenose whales (*Hyperoodon ampullatus*), interacting with benthic trawlers fishing off the eastern Grand Banks of the western North Atlantic in 2007. Whale interactions were only observed when vessels were targeting Greenland halibut (*Reinnhardtius hippoglossoides*) in deep-water fishing areas and were most common during net hauling. Sperm whales and northern bottlenose whales appeared to engage in feeding behavior close to the surface during hauling, especially during the latter stages, suggesting they targeted fish escapees rather than discards. Using photo-identification methods, seven individual sperm whales were identified with multiple resights of six individuals being recorded over an almost two month period. The maximum distance between two resights was 234 km, suggesting individual sperm whales were repeatedly targeting and even following fishing vessels over multiple days and between fishing areas. By contrast, there were no photographic resights of individual northern bottlenose whales within this study, or with substantial photo-identification catalogues from other adjacent high density areas, suggesting that individuals of this species may be less likely to follow vessels or move between areas. This study documents the earliest confirmed records of northern bottlenose whales in this remote region. These interactions and high encounter rates may indicate that adjacent populations are recovering from the previous century of commercial whaling. Our study provides new insights and details on whale-fisheries interactions, which can inform future research and help managers understand the real and perceived impacts on fisheries and whales.

## 1. Introduction

Fishing causes some of the most significant anthropogenic impacts on marine ecosystems [1]. Interactions between cetaceans and fisheries is one major conservation issue and the occurrence of cetacean bycatch, entanglement and depredation of commercial and small-scale fisheries have been reported worldwide [2, 3]. Overfishing can also indirectly threaten cetaceans by decreasing the availability of their prey [4]. However, individual cetaceans may also experience short term benefits from fishing activities if the quantity and the quality of the prey consumed increases and or foraging costs decrease [5]. Bottom trawling accounts for 20% of global fisheries [6] and interactions with bottom trawlers have been observed in at least 19 species of odontocete cetaceans, including sperm whales *(Physeter macrocephalus)* and northern bottlenose whales *(Hyperoodon ampullatus)* [7].

In the western North Atlantic, deep diving sperm whales and northern bottlenose whales have been reported to interact with trawlers targeting species such as Greenland halibut (*Reinnhardtius hippoglossoides*) [8, 9]. One consequence of the recently implemented Import Provisions of the United States *Marine Mammal Protection Act*, is that an improved understanding of the potential for marine mammal bycatch has become of increased importance for fisheries managers [10]. Additionally, research on the risks of fisheries to the Scotian Shelf population of northern bottlenose whales (which have a small population size are considered Endangered status under Canada’s *Species At Risk Act (SARA 2006)* requiring and the development, where necessary, of mitigation measures) has been identified as a management priority.[11]

Responsible ecosystem-based management (EBM) of fisheries requires detailed knowledge of interactions between fisheries and non-target species, including cetaceans. For EBM strategies to be effective, managers need to understand real and perceived impacts on fisheries and on whales. This requires identifying the species involved, the fisheries and marine areas where interactions are most prevalent, and any long-term trends in interactions. In this study we describe sperm and northern bottlenose whales’ interactions with a trawler fishing in the western North Atlantic in 2007 to increase understanding of the characteristics, prevalence, and risks of this associative behavior. Our specific aims are to: (1) identify areas where interactions between fishing activities and sperm and bottlenose whales occur, (2) document the patterns when whales are observed in the fishing cycle and, (3) describe the behaviour of the animals involved.

## 2. Material and Methods

### 2.1 Study area

The study area is located between the western and southern margins of the Grand Banks of Newfoundland in the western North Atlantic (**Figure 1**). This area is managed under the provisions of the Northwest Atlantic Fisheries Organization (NAFO), which requires vessels to carry a fisheries observer on board. The trawler operated in four NAFO divisions: 3L, 3M, 3N and 3O. Fishermen have vernacular names for their preferred fishing areas, where they typically target different species **(Table 1)**. The location of these areas within the NAFO blocks are also shown in **Figure 1**. Fishing effort in each area was decided by the skipper. Trawl tows were typically made parallel to depth contours. The duration of each haul depended on factors such as the nature of the sea bed (depth and relief) and varied between fishing areas **(Table 1)**. Other than towing depth, there was no significant variation in the method of fishing between the main target species (Greenland halibut, redfish (*Sebastes sp*.), or thorny skate (*Raja sp*.)).

**Table 1.** Target species and summary information for each fishing area. Target species varied between areas and included: Greenland halibut (*Reinhardtius hippoglossoides*), Thorny skate (*Raja sp*.*), and* Redfish (*Sebastes* sp). Other species such as grenadiers (*Macrourus spp*), American plaice (*Hippoglossoides platessoides*), witch flounder (*Glyptocephalus cynoglossus*), yellowtail (*Limanda ferruginea*), white hake (*Urophycis tenuis*) and cod (*Gadus morhua*) were also taken.

| NAFO \| Fishing Area |  | Target Species | Depth Range (m) | Tow Cable Length (m) | Towed Distance (m) | Mean Haul Duration (h) | Average Haul Length (km) | Hauls Observed (N) | Observation Periods (N) | Observation Time (h) |
| --- | --- | --- | --- | --- | --- | --- | --- | --- | --- | --- |
| 3L | La Décima | Greenland halibut | 750-1400 | 2500-2700 | 2377±127 | 7.3±1.2 | 36 | 10 | 16 | 12.9 |
|  | Playa Nova |  | 750-1450 | 2200-2700 | 2264±186 | 10.6±1.9 | 59 | 48 | 101 | 90 |
| 3N | O Pequeno | Greenland halibut | 500-1450 | 1700-2800 | 2301±332 | 55.4±1.3 | 46 | 21 | 39 | 31.2 |
|  | Carson Canyon |  | 700-1300 | 2000-2400 | 1957±118 | 8.4±2.4 | 25 | 12 | 27 | 24.9 |
|  | El Seco | Thorny skate | 50- 90 | 340 | 331±4 | 4.9±1.6 | - | 25 | 26 | 21 |
| 3O | Raiz Cuadrada | Greenland halibut | 900- 1100 | 2500 | 2281±60 | 11.6±4.9 | 32 | 2 | 2 | 10 |
|  | Divisoria | Redfish | 550 | 600 | 1775 | 11 |  | 1 | 4 | 0.6 |
| 3M | Flemish Cap | Redfish | 50-800 | 850 | 797±11 | 4.5±1.9 | 18 | 9 | 15 | 11.6 |
| - | Transiting | NA | - | - | - | - | - | - | 16 | 14.3 |
| Total |  |  |  |  |  |  |  | 128 | 248 | 202.2 |

**Figure 1.**
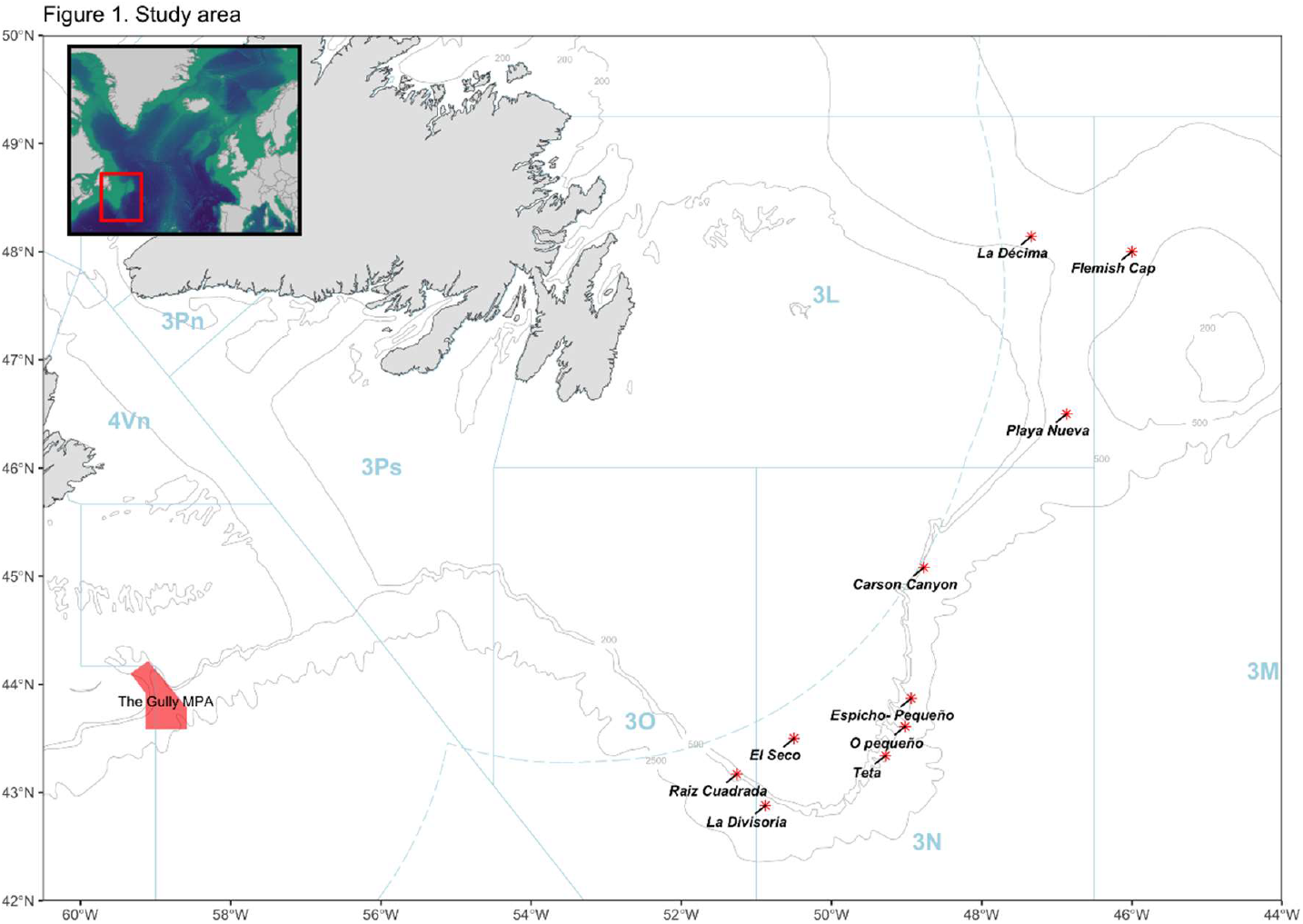
Map of the NAFO convention area, showing NAFO divisions, fishing areas named by fishermen and the Gully marine protected area (MPA).

The study area is located between the two sub-populations of northern bottlenose whales in western North Atlantic recognized by Canada. The Scotian Shelf population inhabits a region off Nova Scotia and the southern Grand Banks of Newfoundland, with high site fidelity to a large submarine canyon called the Gully [12]. Another concentration of northern bottlenose whales can be found in the Davis Strait and Baffin Bay area; however, the size of this population is unknown [13].

Most sightings of sperm whales in the North Atlantic are of single individual males, or more rarely small groups of males. Higher numbers of sperm whales are found along the shelf edge, with occasional sightings in shallower regions [14].

### 2.2 Field observations

Observations on whale encounters were collected on 50 days, between the 20th of July and the 13th of September in 2007, by UO while working as a NAFO observer on the 51 m stern trawler *Playa Menduiña Dos*. Data collection, analysis and publication of this study was conducted with the vessel captain’s consent. Observations of whales were typically made from the vessel’s bridge during the day, with some behavioral observations occurring at night when whales were within 30 m of the boat. Nighttime observations relied on the vessel’s lights, which were directed astern to allow the cod end to be monitored as the net was hauled to the surface.

Data collection was carried out during the four types of vessel operations (**Table 2**). The distance between the trawler and the net varied depending on the fishing area and depth. Trawler speed varied depending on fishing activity (**Table 1**). Information recorded during each observation period included the trawler location, speed, trawling activity, cloud cover, water temperature, sea state, and visibility. Although effort directed at recording whales had to be scheduled around NAFO observer duties, standard data collection protocols were followed, and efforts were made to distribute observation periods across all fishing activities.

**Table 2.** Description of trawler activity states and opportunistic prey availability during each activity.

| Trawler activity | Activity Description | Opportunistic Prey Available |
| --- | --- | --- |
| Preparing/<br>shooting<br>the net | After hauling, the fishermen quickly inspect the net for damage. Some of the crew start processing the fish, releasing bycatch and discarding offal. Once the net has passed inspection, the other crew begin the next net deployment, a process that can take 1- 1.5 hours. During the initial process of deploying the fishing gear to the seabed, the trawler is stopped or moving at very low speed. Shooting commences with the cod end being launched over the stern, then the net is let out as the vessel steams slowly ahead. The trawl | Discards of bycatch and offal from target species |
|  | doors are attached to the gear and the warps slacked away from the winch until the gear is in position on the seabed. The trawl doors provide the horizontal spread of the net. The trawler moves slowly ahead and once the trawl doors are in the water, the rest of the crew proceed to process the remaining fish. Shooting ends when the net reaches the seabed, which can vary based on depth. |  |
| Towing | The action of trawling the fishing gear along the seabed. During towing the crew are still processing the fish. | • Discards of bycatch and offal from target species as catch from previous trawl processed.<br>• Escapees from the net |
| Hauling | The process of retrieving the fishing gear after towing. The vessel trawl winches haul the net and lift the cod-end containing the catch on board from the stern over the deck. The winches generate an intense noise and vibration in the hull when working. | • Escapees from the net |
| Transiting | Navigating to other fishing areas. | • None |

### 2.3 Sperm and northern bottlenose whale data collection

Data on both sperm and northern bottlenose whales were collected during observation periods. Each period included only one type of trawler activity and lasted for approximately 30 minutes. Two field protocols were followed: (1) scan sampling for whale presence and behaviour every ten minutes and (2) recording the duration of whale encounters. An observed encounter was defined as a near continuous observation of a whale at the surface and the encounter was ended when whales were absent (dove or left the area) for ten minutes or more. During a scan sample an entire 360° sector around the trawler was scanned visually for the presence of whales. Data was recorded on whale species and group size and location relative to the vessel. Whale behaviors (**Table 3, Figure 2 and 3**) were recorded when observation conditions allowed (animals within 300 m, sea state < Beaufort 4).

**Table 3.** Whale behaviour definitions. SW = sperm whale, NBW = northern bottlenose whale.

| Whale behavior | Definition | Species |
| --- | --- | --- |
| Swim parallel | Swimming alongside and at the same speed as the trawler. | SW |
| Following | Swimming astern the trawler at a constant speed and distance (**Figure 2**). | SW, NBW |
| Swimming in an opposite direction | Swimming with no apparent association with the heading of the trawler. | **SW, NBW** |
| Fluke up | Tail fluke is raised high into the air, initiating a steep dive. | **SW** |
| Surface feeding | Typically observed during the final stage of hauling when the cod end was close to the vessel's stern and fish were observed escaping the net as seabirds (e.g., great shearwaters; *Ardenna gravis*, northern fulmars; *Fulmarus glacialis*) brought them to the surface. | **SW, NBW** |
|  | Slowly turns body sideways or upside down just below the surface, showing first the dorsal fin, then the blowhole, and finally, their whitish jaw. | **SW** |
|  | Makes repeated short shallow dives, during surfacing, with their beak frequently visible | **NBW** |
| Breach | A whale lifting much of its body above the water before splashing back down.(**Figure 3**). | **SW, NBW** |
| Lobtail | Energetic tail slaps at the surface. | **SW, NBW** |

**Figure 2.**
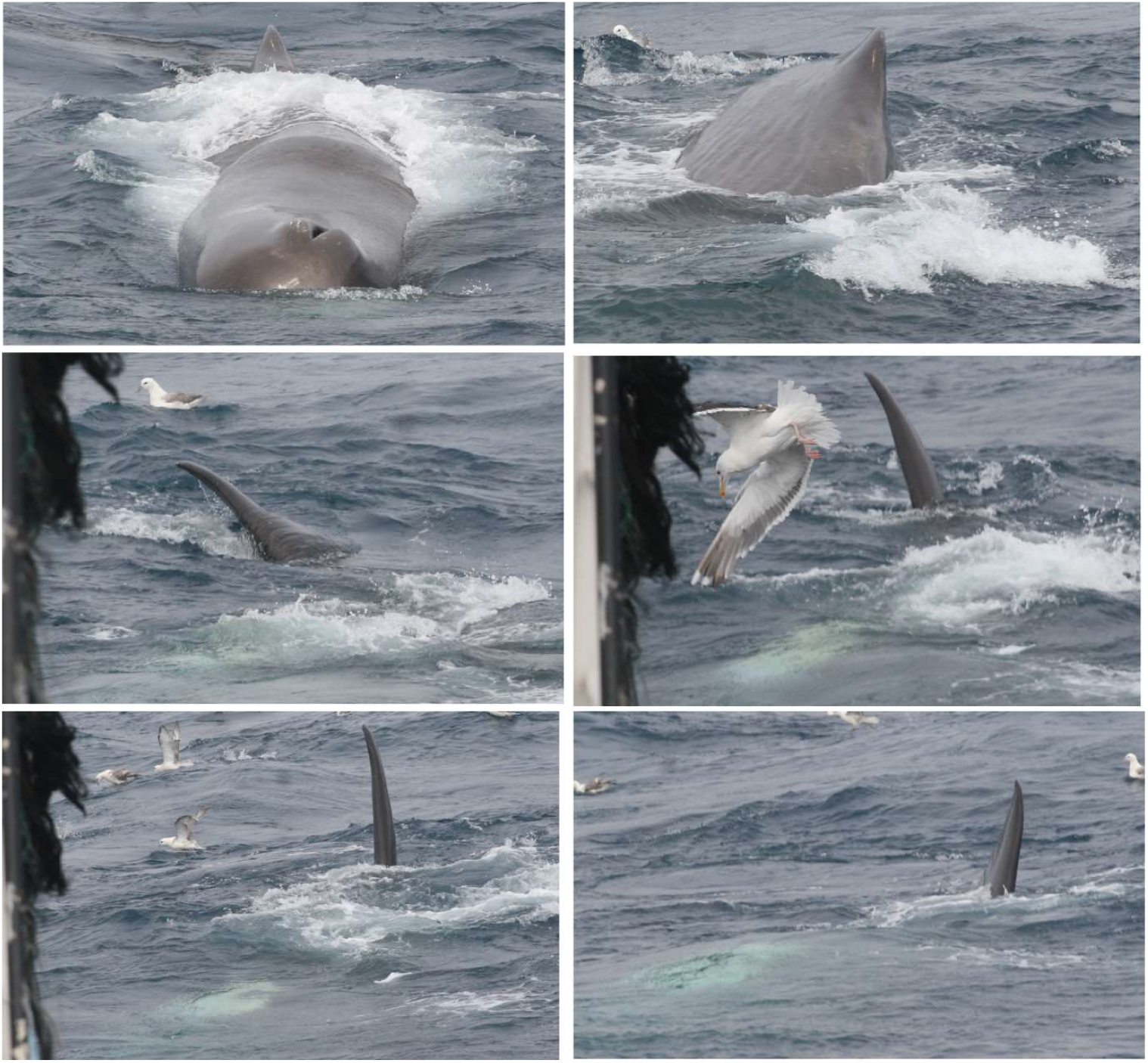
Sequence showing a sperm whale turning his body to the right. First picture shows blowhole and dorsal fin, second turning to the right, dorsal fin and part of the side is observed, in the last four pictures the fluke is going to the right. In the last picture the sperm whale is almost upside down. We associated this pattern of observations with surface feeding close to the surface.

**Figure 3.**
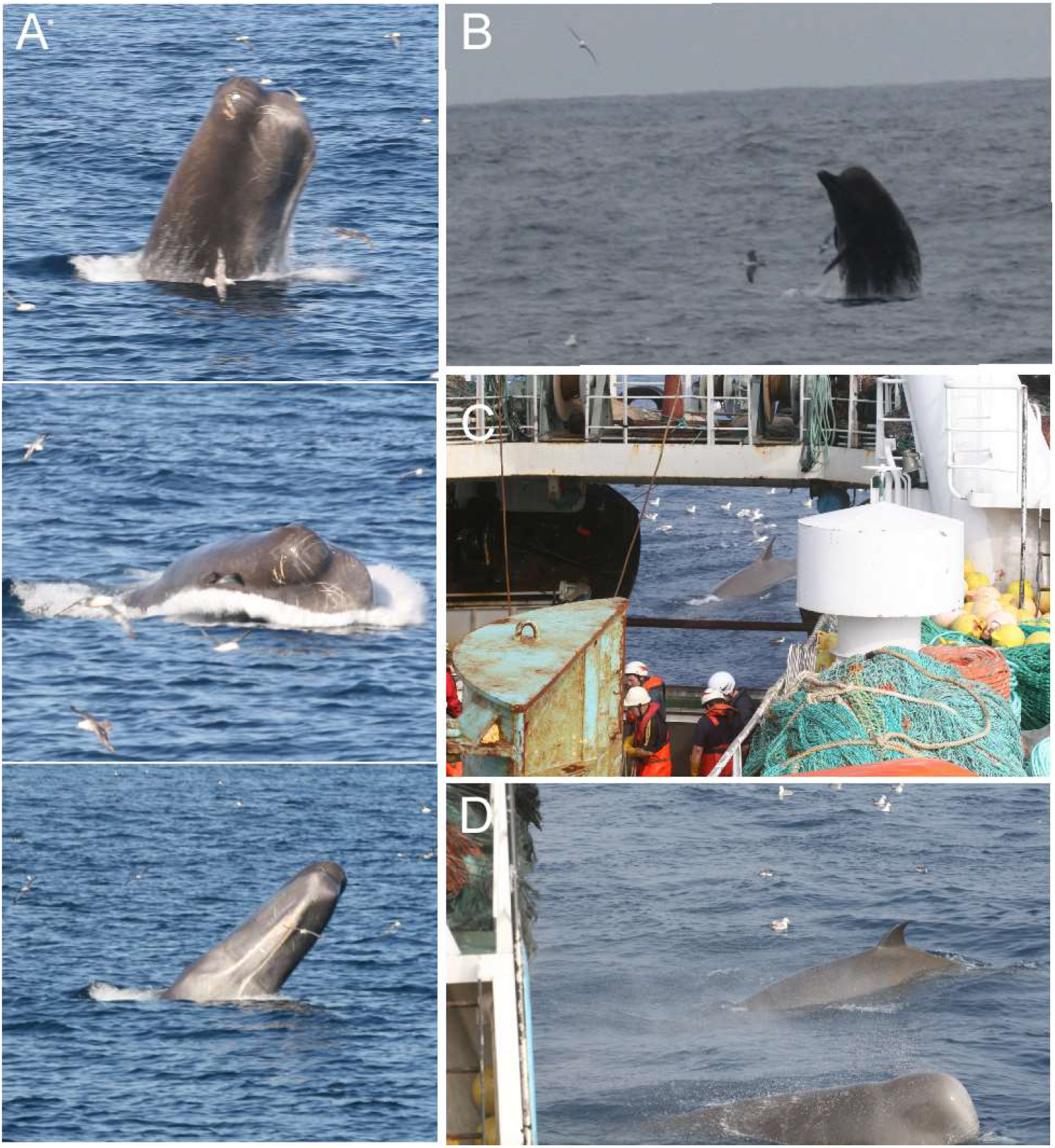
Sperm whale (A) and northern bottlenose whale (B) breaching. Northern bottlenose whales close to the boat at the end of hauling (C-D).

When possible, identification photographs were taken of the dorsal fin and the surrounding flank area of both species, and of the flukes of sperm whales using a digital camera (Canon EOS 30) with a 200 mm lens. Photographs were taken irrespective of the presence of any obvious marking on the individual, and efforts were made to obtain multiple photographs of every individual in each encounter.

### 2.4 Photo-identification analysis

Photographs were used to identify individuals, as well as confirm group size and species. Individual whales can be reliably identified from various identifying nicks, scars and marks, typically on the trailing edge of the flukes (sperm whales) and on the melon, back and dorsal fin (sperm whales and northern bottlenose whales) [15, 16]. In this study identifiable marks that were visible anywhere on an individual’s body (i.e., backs, dorsal fins, melons, flukes) were used to distinguish individual whales within this relatively short study. Matching with northern bottlenose whale photographic catalogues external to this study only considered marks that are known to be persistent and reliable for long-term identification (e.g., notches and back indents) [17]. Images of northern bottlenose whales were quality (Q) using established criteria for photographic identification (Feyrer et al. 2021) and only images of Q3 or above were included in matching between catalogues.

Associations between individual sperm whales were defined by co-occurrence within the same observation period as defined above in section 2.3. Where the same photographic perspective on an identifying mark could be compared (e.g., tail fluke), individuals were identified across encounters. Sperm whale fluke photographs were added to Flukebook which used computer matching algorithms to compare images with those submitted by other researchers in the North Atlantic [18].

The average observed encounter rates were calculated by dividing the number of encounters by the length of each observation period. The average observed encounter rate was calculated for each trawler activity state and each area. To assess the relationship(s) between different fishing areas and trawler activity state on whale presence we used binomial generalized linear models (GLMs). The response variable was presence or absence of whales during observation periods with model terms for fishing area, trawler activity state, sighting conditions (i.e., Beaufort Sea state, visibility (m), cloud cover (%) and daylight), and an offset for effort (length of observation time). Models for sperm whales and northern bottlenose whales were built separately. Observations made in fishing areas with < 10 observation periods, were excluded to avoid model separation due to singularities. Transiting between areas was not included in models as there was no corresponding region associated with the activity. Dispersion and zero inflation of all GLMs were checked with the DHARMa package (Hartig, 2017), which approximates dispersion and zero inflation via simulations.

The ‘fluke-up’ behaviour of sperm whales, during which their entire tail fluke emerges from the water, indicates an individual was initiating a steep dive (probably related to foraging). The mean fluke up rate was calculated by dividing the number of sperm whale ‘fluke ups’ recorded in an observational unit by the total time for that unit. Rates were compared for each trawler activity state, using Kruskal Wallis and Tukey test. All statistical analyses were completed in R [19].

## 3. Results

Overall, 200 hours of fishing activities were monitored for whales across 50 days. The majority of the observation effort was evenly distributed between the hours of 8:00 - 12:00 am, with 8.3 hours of observation occurring at night (between 21:30 and 6:30). A total of 248 observational periods were completed in seven different fishing areas and while transiting between sites **(Table 4)**. Observational effort occurred during 104 (81.25%) of the 128 net deployments made by the vessel during the study period. The total hours of observation, trawler speed and whale behavior during each trawling activity is summarized in **Table 5**. Summaries of the hauls monitored in each fishing area (i.e., number, average duration, observational effort, net distance and depth, total days and target species) are shown in **Table 1**.

**Table 4.** Fishing areas with summary information for encounters with sperm whales and northern bottlenose whales. Encounter rates are calculated per observation period and averaged for all observations in an area, calculated with a 95% bootstrap confidence interval. Differences in summary totals due to time spent transiting between areas and values for all trawler states are included in Table 5.

|  | *Carson*<br>*Canyon* | *La*<br>*Décima* | *Raiz*<br>*Cuadrada\** | *O*<br>*Pequeno* | *Playa*<br>*Nova* | *Divisoria* | *El*<br>*Seco* | *Flemish*<br>*Cap\** | **Mean**<br>**(95%**<br>**CI)** | **Total** |
| --- | --- | --- | --- | --- | --- | --- | --- | --- | --- | --- |
| Total time observed (h) | 23.2 | 12.3 | 1.8 | 28.8 | 82.6 | 7.6 | 18.2 | 10.7 | 22.2 | 185.2 |
| Observational units | 27 | 16 | 2 | 39 | 101 | 5 | 26 | 15 | 28.8 | 213 |
| Number encounters SW | 22 | 19 | 6 | 13 | 85 | 0 | 0 | 1 |  | 148 |
| Number encounters NBW | 2 | 11 | 1 | 22 | 59 | 2 | 0 | 1 |  | 98 |
| Number encounters both<br>species at same time<br>(included in species total) | 2 | 4 | 1 | 6 | 22 | 0 | 0 | 1 |  | 36 |
| Sperm whale encounter<br>rate (h<sup>-1</sup>) ±SE | 0.42±<br>0.12 | 1.08±<br>0.18 | 1.57± 0.82 | 0.58±<br>0.18 | 0.63±<br>0.20 | 0.0± 0 | 0.0 | 0.10±<br>0.10 |  | - |
| NBW encounter rate (h<sup>-1</sup>)<br>±SE | 0.10±<br>0.07 | 0.63±<br>0.20 | 0.37± 0.38 | 0.78±0.2<br>1 | 0.58±<br>0.09 | 0.30±<br>0.30 | 0.0 | 0.10±<br>0.10 |  | - |
| Max number SW<br>observed in an encounter | 2 | 1 | 3 | 2 | 6 | - | - | - | 2.8 | - |
| Mean number SW<br>observed per encounter | 1.4±0.5 | 2.1±1.1 | 1.5±0.8 | 1.1±0.3 | 1.6±0.9 | - | - | - | 1.5±0.4 | - |
| Max number BW<br>observed in an encounter | 5 | 9 | 2 | 5 | 5 | 1 | - | - | 4.5 | - |
| Mean number BW<br>observed per encounter | 3±0 | 2.5±2.3 | 3±0 | 2.5±1 | 2.7±1.2 | 1 | - | - | 2.3±0.7 | - |

\*Only one encounter in these fishing areas

**Table 5.** Fishing trawler activities and summary information for encounters with sperm whales and northern bottlenose whales. Encounter rates are calculated per observation period and averaged for all observations in a trawler activity state, calculated with a 95% bootstrap confidence interval.

|  | Preparing/<br>Shooting | Towing | Hauling | Transiting | Summary Total |
| --- | --- | --- | --- | --- | --- |
| Mean speed (knots) ± SD | 5.37±2.57 | 3.04±0.5<br>2 | 1.44±0.89 | 6.78±2.57 | - |
| Observational units | 68 | 82 | 82 | 16 | 248 |
| Total time observed (h) | 48.9 | 85.0 | 51.6 | 14.3 | 199.8 |
| No. encounters SW | 24 | 24 | 44 | 2 | 129 |
| No. encounters NBW | 20 | 15 | 46 | 5 | 86 |
| No. encounters both species<br>(Included in species total) | 10 | 2 | 21 | 1 | 34 |
| Sperm whale encounter rate ± SE (h<sup>-1</sup>) | 0.53± 0.11 | 0.35<br>±0.07 | 0.67± 0.09 | 0.19± 0.13 | - |
| NBW encounter rate (h<sup>-1</sup>) | 0.45± 0.10 | 0.22<br>±0.08 | 0.73± 0.11 | 0.40± 0.16 | - |
| Total no. of flukes | 4 | 14 | 26 | 0 | 44 |
| Average fluke rate (h<sup>-1</sup>) | 0.1 | 0.2 | 0.5 | 0 | - |
| Max no. SW observed in an encounter | 6 | 5 | 5 | 1 | - |
| Mean no. SW observed per encounter<br>± SD | 1.8±1.1 | 1.2±0.4 | 1.8±0.9 | 1 | - |
| Max no. NBW observed in an<br>encounter | 5 | 9 | 5 | 3 | - |
| Mean no. NBW observed per<br>encounter ± SD | 2.7±1.1 | 2.0±1.9 | 2.8±1.2 | 2.3±0.5 | - |

### 3.1 Whale sightings

We observed four species of cetaceans: common dolphins (*Delphinus delphis*, N = 6 encounters), long-finned pilot whales (*Globicephala melas*, N = 3 encounters), sperm whales (N=129 encounters) and northern bottlenose whales (N=86 encounters). Northern bottlenose whales and sperm whales were the only species observed interacting with the trawl. The two species were seen together on 34 occasions with no apparent inter-species interaction **(Figure 4)**.

**Figure 4.**
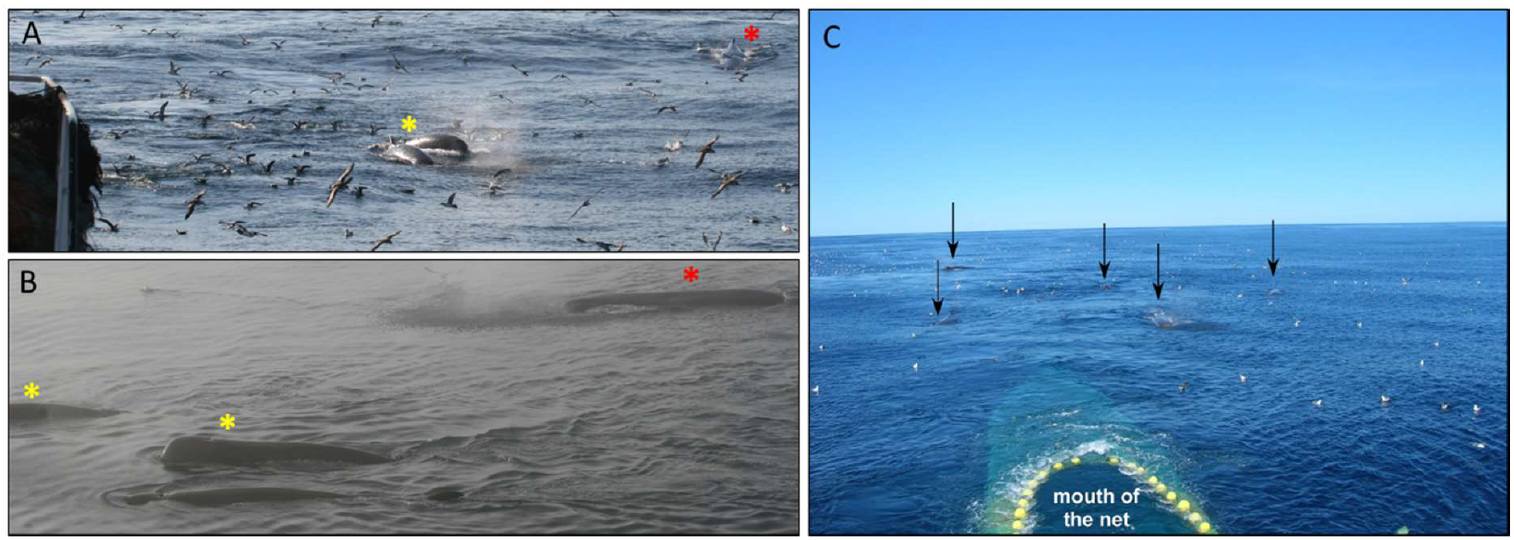
Examples of whale behaviors. **(A)** and **(B)** Sperm and northern bottlenose whales seen together. **(C)** Five sperm whales following the trawler at the end of hauling with the mouth of the net visible at the surface behind the yellow buoys. Red stars and black arrows indicate sperm whales and yellow stars indicate northern bottlenose whales.

Observations were made during 14 nighttime hauls. There were were four encounters with sperm whales and six encounters with northern bottlenose whales during these. Whale behavior at nighttime was similar to that observed during hauling in the daytime (i.e., surface feeding, fluke ups).

Datasets used for modelling trends in observed encounters are available as part of the open access data repository for this publication [20].

#### 3.1.1 Whale presence across fishing areas

Both sperm and bottlenose whales were observed in four fishing areas: *La Décima* (3L), *Playa Nova* (3L), *Carson Canyon* (3N) and *O Pequeno* (3N), while northern bottlenose whales were also sighted in *Raíz Cuadrada* (3O). These areas are generally on or close to the slope edge, where depths are between 700-1450. Greenland halibut was the target species in these areas. Neither species was sighted in *Flemish Cap* (3M) nor *El Seco* (3N), where fishery targets were redfish and thorny skates respectively and water depths were much shallower 50-90m (*El Seco*), and 50-800m *(Flemish Cap*) **(Figure 5)**.

**Figure 5.**
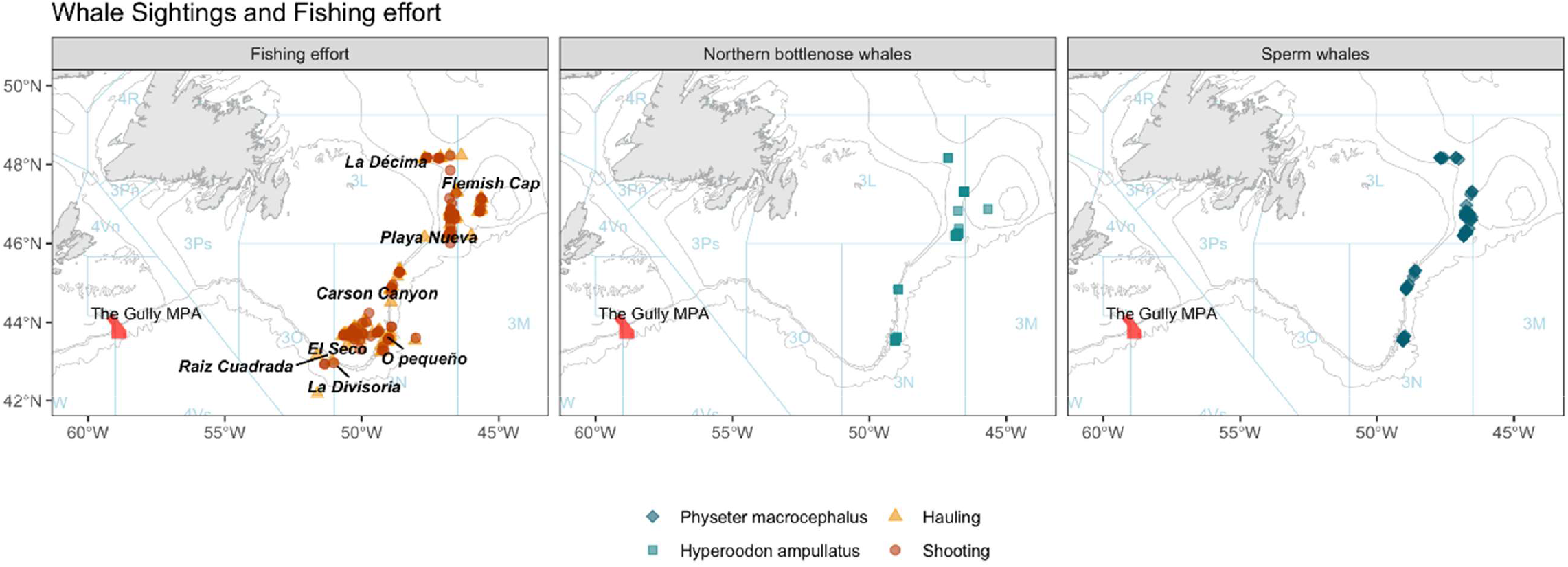
Location of hauling and shooting effort in relation to sightings of sperm whales and northern bottlenose whales.

Across this study, an average of 1.5 sperm whales were sighted during an observed encounter. *La Décima* had the highest average observed sperm whale encounter rate of 1.08/hr. of observation, while the largest group size of sperm whales (N = 6 individuals) was observed in *Playa Nova*. For northern bottlenose whales, observed encounter rate was highest (0.78/hr. of observation) in *O Pequeno*. The maximum number of northern bottlenose whales in an observed encounter was 9, in *La Décima*, and the mean number of individuals across all observed encounters was 2.3 (**Table 4, Figure 6**).

**Figure 6.**
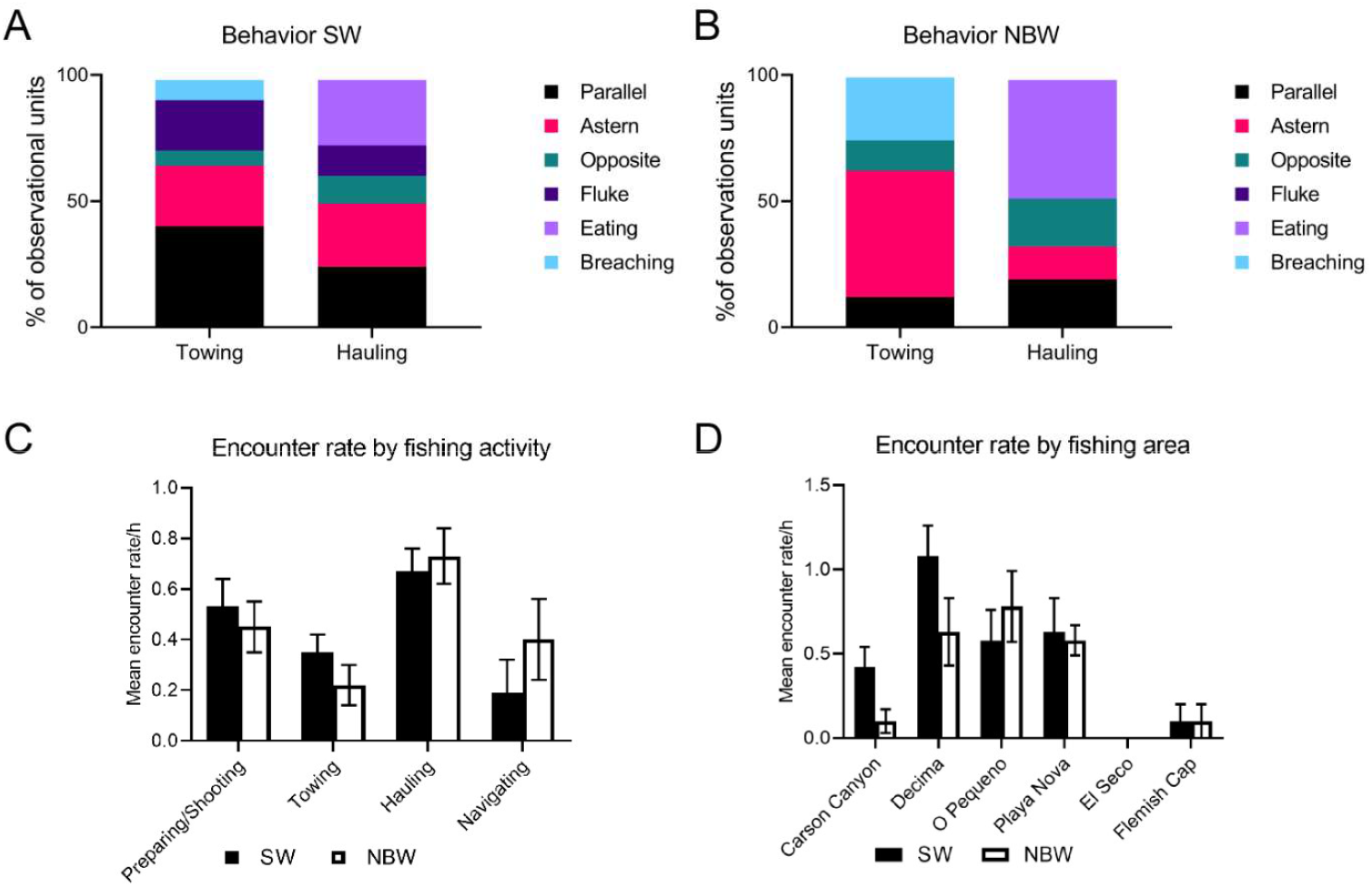
The occurrence of the different whale behaviours versus fishing area and trawler activity. **(A)** Sperm whales **(B)** Nothern bottlenose whales’ behavior per trawling activity. Mean whale encounter rates per **(C**) trawling activity and **(D)** fishing area. Six behaviors were scored: (1) swimming parallel to the vessel, (2) following the trawler at a constant speed and near constant distance astern, (3) swimming in the? opposite direction?? from the trawler (normally within 50 m), (4) fluke, (5) surface feeding, and (6) breaching/lobtailing.

#### 3.1.2 Whale behavior and trawler activity states

The behavior of both sperm and northern bottlenose whales differed with trawler activity, particularly between towing and hauling (**Table 5**). During towing sperm whales were often observed swimming parallel to and either abeam or astern of the trawler while maintaining a constant speed and distance **(Figure 4)**. The longest sighting of a sperm whale, one hour and 25 minutes of continuous observation, occurred during towing. Surface feeding behavior was not observed during towing but lobtailing and repeated breaching was observed on one occasion (**Figure 3)**. Northern bottlenose whales disappeared soon after shooting the net but were often observed swimming behind the net as it was being hauled. On two occasions repeated breaching was observed while towing (**Figure 3B**). Feeding behavior on discards was not observed (**Table 2**).

During hauling, the boat slows down, and the net winches start hauling back the net, generating a loud noise and vibration in the hull. At this time, both sperm whales and northern bottlenose whales were observed astern, swimming towards the trawler, or surface feeding. Sperm whale “fluke ups” were observed five times more frequently during hauling, than preparing/shooting, and ∼2.5 times more often than during towing (Kruskal-Wallis test, Tukey HSD p<0.05, N=248). The fluke-up rate per hour was significantly higher during hauling (0.5 fluke-ups per whale) than during preparing/shooting and towing (Kruskal-Wallis test, Tukey HSD p<0.05, N=248) (**Table 5**). Surface feeding behavior was only observed at the end of hauling, when the cod end was at or close to the surface, and coincided with birds feeding on escaping fish at the surface. Overall, observed encounter rates for both sperm whales and northern bottlenose whales were significant higher during hauling than during all other activity states (**Figure 6C**).

#### 3.1.3 Whale presence, fishing area and trawler activity states

Fishing area and trawler activity state were both significant terms in modelling northern bottlenose whales and sperm whale presence. Sighting conditions were also significant in predicting observations of both species; however, the affecting sperm whale sighting rates variables (i.e., sea state, visibility, daylight), were different from those which were significant for northern bottlenose whales (i.e., cloud cover, daylight). GLMs with these variables had significantly more support than models with additional or fewer terms (**Table 6 A, B**). Inspection of coefficients indicated that *La Décima* and *Playa Nova* were significant areas (p<0.05) relative to *Flemish Cap* for the presence of both sperm whales and northern bottlenose whales. Two areas, *Raiz Cuadrada* and *Divisoria*, were not included in the models due to low observation effort (**Table 1**). All trawler activities — Towing (T), Hauling (H), and Preparing or Shooting the Net (P/S), were significantly associated with sperm and northern bottlenose whale presence (**Table 6**).

**Table 6.**
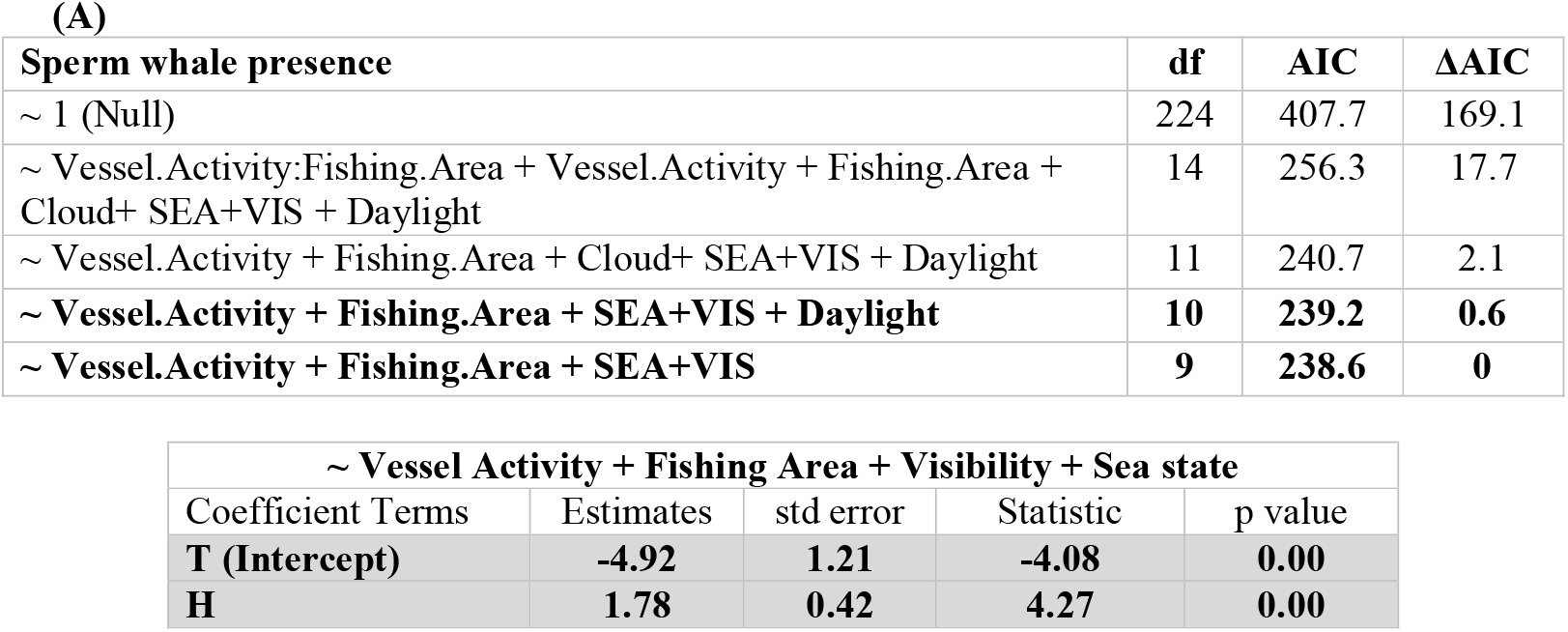
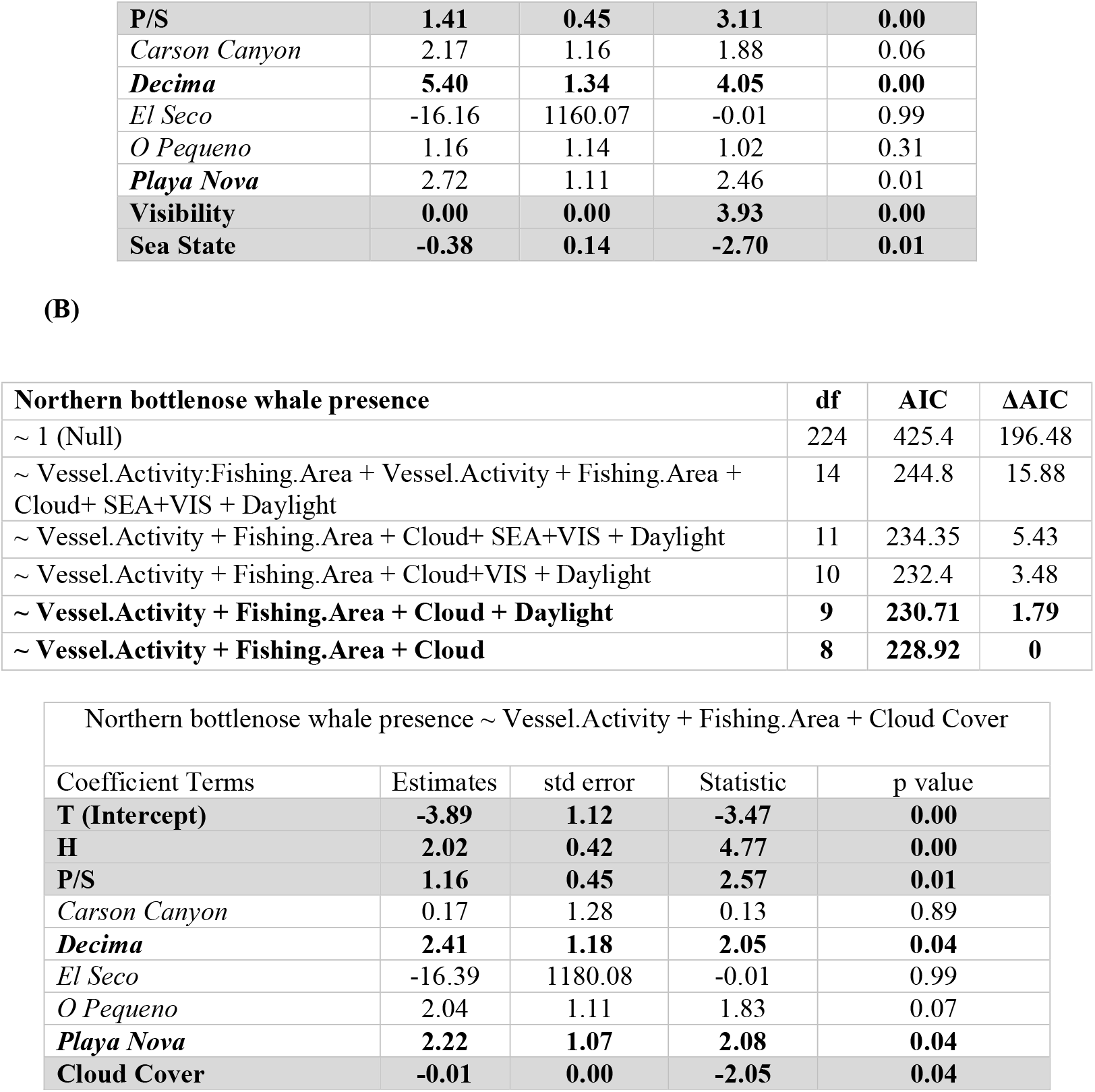
Summary of binomial generalized linear regression models (GLMs) used to assess the effect of trawling activity, region and sighting conditions on observations of (A) sperm whales or (B) northern bottlenose whales. Greatest support for best fit models is indicated by lowest ΔAIC (Akaike’s information criterion) values; all model with ΔAIC < 2 indicated in bold. Coefficients from the most supported model are provided, and terms where support for a significant relationship (positive or negative) with whale presence (p-value <0.05) are noted in bold. For Trawler Activity state, Towing was the reference level and for Fishing Area, Flemish cap was the reference level.

| Sperm whale presence | df | AIC | $\Delta AIC$ |
| --- | --- | --- | --- |
| ~ 1 (Null) | 224 | 407.7 | 169.1 |
| ~ Vessel.Activity:Fishing.Area + Vessel.Activity + Fishing.Area + Cloud+ SEA+VIS + Daylight | 14 | 256.3 | 17.7 |
| ~ Vessel.Activity + Fishing.Area + Cloud+ SEA+VIS + Daylight | 11 | 240.7 | 2.1 |
| ~ **Vessel.Activity + Fishing.Area + SEA+VIS + Daylight** | **10** | **239.2** | **0.6** |
| ~ **Vessel.Activity + Fishing.Area + SEA+VIS** | **9** | **238.6** | **0** |

| ~ Vessel Activity + Fishing Area + Visibility + Sea state |  |  |  |  |
| --- | --- | --- | --- | --- |
| Coefficient Terms | Estimates | std error | Statistic | p value |
| **T (Intercept)** | **-4.92** | **1.21** | **-4.08** | **0.00** |
| **H** | **1.78** | **0.42** | **4.27** | **0.00** |
| **P/S** | **1.41** | **0.45** | **3.11** | **0.00** |
| *Carson Canyon* | 2.17 | 1.16 | 1.88 | 0.06 |
| ***Decima*** | **5.40** | **1.34** | **4.05** | **0.00** |
| *El Seco* | -16.16 | 1160.07 | -0.01 | 0.99 |
| *O Pequeno* | 1.16 | 1.14 | 1.02 | 0.31 |
| ***Playa Nova*** | **2.72** | **1.11** | **2.46** | **0.01** |
| **Visibility** | **0.00** | **0.00** | **3.93** | **0.00** |
| **Sea State** | **-0.38** | **0.14** | **-2.70** | **0.01** |
| (B) |  |  |  |
| **Northern bottlenose whale presence** | **df** | **AIC** | **ΔAIC** |
| ~ 1 (Null) | 224 | 425.4 | 196.48 |
| ~ Vessel.Activity:Fishing.Area + Vessel.Activity + Fishing.Area + Cloud+ SEA+VIS + Daylight | 14 | 244.8 | 15.88 |
| ~ Vessel.Activity + Fishing.Area + Cloud+ SEA+VIS + Daylight | 11 | 234.35 | 5.43 |
| ~ Vessel.Activity + Fishing.Area + Cloud+VIS + Daylight | 10 | 232.4 | 3.48 |
| **~ Vessel.Activity + Fishing.Area + Cloud + Daylight** | **9** | **230.71** | **1.79** |
| **~ Vessel.Activity + Fishing.Area + Cloud** | **8** | **228.92** | **0** |

| Northern bottlenose whale presence ~ Vessel.Activity + Fishing.Area + Cloud Cover |  |  |  |  |
| --- | --- | --- | --- | --- |
| Coefficient Terms | Estimates | std error | Statistic | p value |
| **T (Intercept)** | **-3.89** | **1.12** | **-3.47** | **0.00** |
| **H** | **2.02** | **0.42** | **4.77** | **0.00** |
| **P/S** | **1.16** | **0.45** | **2.57** | **0.01** |
| *Carson Canyon* | 0.17 | 1.28 | 0.13 | 0.89 |
| ***Decima*** | **2.41** | **1.18** | **2.05** | **0.04** |
| *El Seco* | -16.39 | 1180.08 | -0.01 | 0.99 |
| *O Pequeno* | 2.04 | 1.11 | 1.83 | 0.07 |
| ***Playa Nova*** | **2.22** | **1.07** | **2.08** | **0.04** |
| **Cloud Cover** | **-0.01** | **0.00** | **-2.05** | **0.04** |

### 3.2 Photo identification analysis

A total of 7,343 photographs were taken during whale encounters for photo-identification analyses. Twenty-three northern bottlenose whale individuals were identified based on photographs of the right-side dorsal fins and 15 individuals were identified from left-side dorsal fin photographs (**Figure 7**). None of the northern bottlenose whales identified were reidentified between hauls or on multiple days.

**Figure 7.**
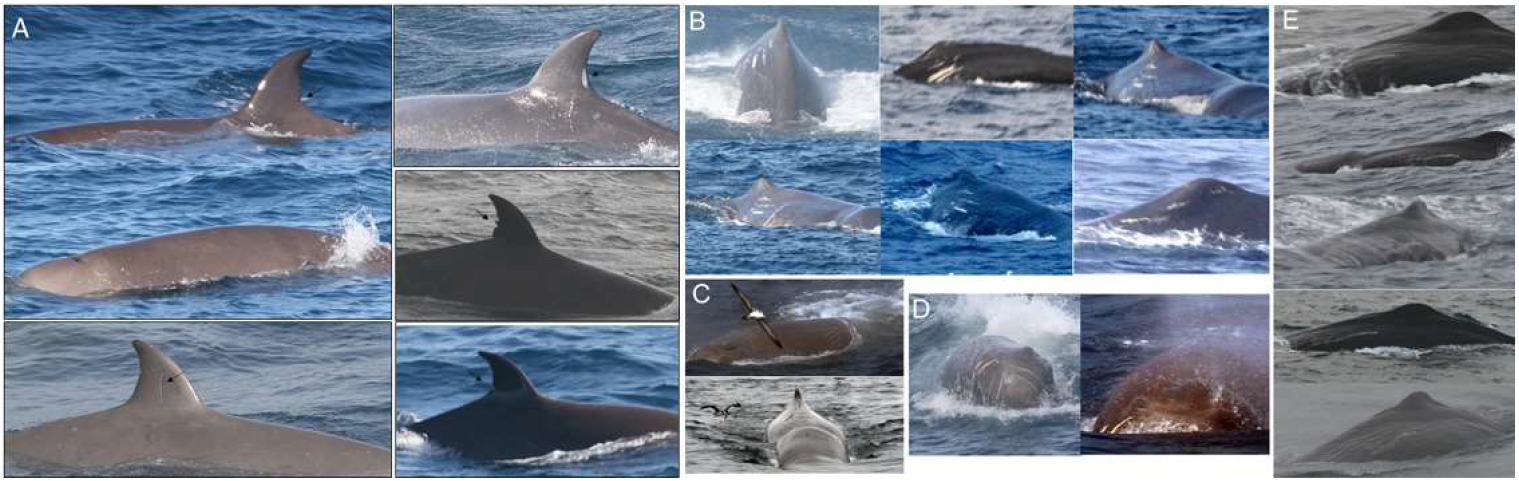
Examples of distinct marks used for. **(A)** bottlenose whale identification, including notches and large fin scars and **(B)** for sperm whale identification. *Neboa*, with characteristic scars at both sides of the dorsal fin; *Scratchy* showed a large lateral scratch; *Breixo*, showed many scars in the front part of his head; and *Faneca*, shows a lateral scar visible only from the left side, but very easy to identify.

A total of seven sperm whales were identified using markings on their tail flukes, with an additional three whales identified solely by distinctive marks on their bodies **(Figure 7 and Table 7**). These ten distinctively marked individuals were named for the observer’s reference and tracking within encounters (**Figure S1, S2**). However, due to the non-overlapping photographic perspective of individuals, duplicate identifications could have occurred across encounters in this study. After this was accounted for, the minimum number of individual sperm associating with the trawler, as determined by distinct markings on the tail fluke was seven. Photographs of sperm whales encountered in this study are publicly available to other researchers through the Flukebook platform.

*Scratchy* was seen on two occasions, 11 days apart, with a distance between encounters of only 4 km. *Neboa* was encountered eight times in a week while fishing in *Playa Nova*, across more than 100 km (**Table 7, Figure S2)**. The calculated minimum average swim speed for *Neboa* over this period was 4.0 km/h. *Sparrow* was observed repeatedly with only 5-16 km between encounters, but then seen again on two consecutive days, 58 km away. *Faneca* was encountered five times on three consecutive days in *Carson Canyon. Breixo*, one of the first whales to be identified, was seen during five encounters in *Playa Nova* and subsequently 20 days later and 235 km away in *Carson Canyon*; the maximum distance between two sightings of an individual sperm whale in this study. There are no indications of preferred associations between individuals within this limited dataset, however we note that *Breixo* was seen with each of the other identified whales **(Fig. S1**).

## 4. Discussion

In this paper we have described the behavioral associations of sperm and northern bottlenose whales with trawling activities in the western North Atlantic, supporting a history of anecdotal observations made by fishermen prior to this study and cetacean researchers in the area in 2016-2017 [21]. Whales were most frequently encountered when the trawler was engaged in hauling while targeting Greenland halibut in deep waters close to the continental shelf break[12, 22].

While we were not able to record the whale’s underwater foraging behavior directly, both species engaged in surface feeding and sperm whales increased their rate of fluke ups during hauling. Suggesting that whales may primarily benefit by feeding on fish escaping from the net during the later stages of hauling.

Associations between whales and fishing vessels are likely to both increase prey encounter rates and to reduce energetic costs associated with greater dive depths [5], which corresponds to our observations of increased surface activity during hauling. The size of fish escaping from nets if affected by the mesh size. (It has been shown that for the 135mm mesh size used I NAFO areas 100% retention doesn’t occur until fish are ∼55cm long [23]. Most net escapees will typically be disoriented or disabled, which would make them easier to detect (i.e., either visually or acoustically) and capture. Foraging opportunities provided by trawl fisheries are likely to explain the number of whales that forage opportunistically in this and other areas [5].

Feeding on discards and offal has been described as the primary foraging behavior described for both sperm and northern bottlenose whales associating with Greenland halibut fisheries in the Arctic (Johnson et al 2020, DFO unpublished data). This did not seem to be the case in this study. Discards are released from the rear port side of the trawler, while catch are being processed and can take several hours to complete. However, whales were not seen near the discard valve nor were observation rates correlated with the timing of discards. This suggests that differences in fishing practices (i.e., over time or between regions) may change whale behavior and affect the nature of associations with fishing activities. Understanding of these differences could provide useful insights for management and risk mitigation for whales that associate with these fisheries.

### Sperm whales

Karpouzli & Leaper’s earlier research on interactions between sperm whales and benthic trawlers in the same area in 1997 also showed a pronounced difference in sighting rates with fishing activity, with encounter rates higher during hauling (0.61 sightings per hour), compared to shooting and towing (∼0.02). Our study also revealed high sighting rates during hauling (0.73 sightings per hour).

During the towing phase the net is 1 to 2 kms behind the boat making observations by an observer based on the boat more difficult. It is difficult therefore to rule out the possibility whales are also feeding around nets during towing. A few observations indicate to us that this unlikely to be happening to any great extent. Sperm whales can be seen at these ranges when conditions are favorable and we did not detect whales behind the boat and above the net nor did we note fluke up behavior in this zone. By contrast, it was quite common to see whales swimming at and close to the surface and heading in the same direction as the vessel. during towing and heading. Hauling was also associated with a significantly higher fluke up rates for individual sperm whales; this distinct behavioral change, suggests that opportunistic foraging increases during hauling.

Whether and how whales may be engaging in foraging during other fishing activities, such as preying on the fish escaping the net during towing requires further research Whale behavior during towing and in particular any interaction with the net could be studied using passive acoustic techniques with towed hydrophones and possibly trawl cameras [24].

Male sperm whales tend to be solitary or occur in small groups at high latitudes, and coordinated behavior involving multiple male sperm whales has only been observed on rare occasions. The mean group size of 1.6 animals observed in this study is higher than has been reported for male sperm whales in other high latitude areas in the North Atlantic (e.g., 1.1 for Whitehead et al. 1992; 1.2 for Weir et al. 2001), but smaller than the group size of 2.2, reported in an earlier study of sperm whale trawler interaction in the same area by Karpouzli & Leaper (2004). In this study, the maximum group size of sperm whales was six. This, and the larger average group sizes reported during trawler associations suggest that the presence of fishing trawlers can facilitate or lead to, larger average group sizes. Although sperm whale associations with fisheries occur in other areas, coordinated behavior involving multiple male sperm whales has rarely been observed [16]. However, a recent study by Kobayashi et al. (2020) describes the long-term associations in male sperm whales in the North Pacific, suggesting that male groups may serve to enhance foraging success or provide protection against predators[25].

Six of the photo-identified sperm whales were resighted during multiple encounters, even though conditions for photo-identification were far from ideal. The maximum distance between two sightings of *Breixo*, 20 days apart, was 234 km. Meanwhile, *Neboa* was seen on three consecutive days, with a series of sightings being spread over more than 100 km. The individual sperm whales identified in this study were resighted on different days and in different areas, suggesting they may have followed the trawler and that some individuals may specialize in associative behavior.

Photographs of sperm whale flukes from this study have been submitted to Flukebook [26], an image analysis and database tool for photoidentification used by sperm whale researchers to understand the movements, social associations and population dynamics of sperm whales worldwide. ID photos taken by fisheries observers have the potential to increase our knowledge of whales in remote and otherwise hard to study regions of the ocean.

### Northern bottlenose whales

There are no other published records of northern bottlenose whales associating with fisheries off Newfoundland prior to 2007, here we document the first behavioral records in an area located between two known population centers in the western North Atlantic. Although northern bottlenose whales are often curious and approach vessels [27, 28], there are very few other reports of interactions between northern bottlenose whales and fishing trawlers in the western North Atlantic. Interestingly, Karpouzli & Leaper’s (1997) study whale interaction study, which took place from 1996-1997 in the same area, did not report any observations of northern bottlenose whales. However, in Fertl and Leatherwood’s (1997) review of whale trawl interactions, they identified 15 records from the Scotian Shelf (from unpublished Fisheries and Oceans Canada data) in which northern bottlenose whales were reported to have “followed a trawl during haulback.” Since then Johnson, *et al* (2020) has reported that interactions between northern bottlenose whale groups and fishing vessels, has been ongoing in the Davis Strait-Baffin Bay region of the Canadian Arctic over the last decade. Although, Feyrer, *et al*. (2021) reported photographic evidence of body scarring indicating a steady rate of entanglement scars around the fins of northern bottlenose whales from the Scotian Shelf between 1988-2019, the fisheries responsible could not be identified. Whether associative behavior in northern bottlenose whales is specific to trawling, to certain individual whales and may be spreading as the western North Atlantic populations recover from small population sizes due to commercial whaling, is unclear and requires further study.

Photoidentification showed no resightings of northern bottlenose whales over the course of this study. Here, the minimum number of unique individuals identified (n = 23) is equivalent to ∼16% of the Scotian Shelf population involved in trawl interactions, based on O’Brien and Whitehead’s (2013) population estimate of ∼143. Together with the sighting rates and mean group size, it suggests there is a significant density of animals in the study area. The lack of resights also suggest that individuals are less likely than sperm whales to follow trawlers between trawling events and perhaps are less likely to specialize in this form of foraging. Northern bottlenose whales on the Scotian Shelf are known to have long term high site fidelity, with low rates of movement between areas [12, 28]. Higher observed encounter rates in *Playa Nova* and *La Décima* could indicate that these too are important areas for northern bottlenose whales.

The study area occurs in the border region between the two sub-populations of northern bottlenose whales in the western North Atlantic - the Scotian Shelf and Davis Strait-Baffin Bay population. A recent genetic assessment of northern bottlenose whale population structure by Feyrer, *et al*. (2019) included samples collected near *La Décima* (i.e., “Newfoundland”) and found this region to be an area of mixing between the Scotian Shelf and Labrador-Davis Strait populations. Our study area, from *La Décima* in the north to *Raíz Cuadrada* in the south, lies mid-way between both populations (Figure 1). A review of the photographic identification catalogues from the Scotian Shelf (1988-2019), Newfoundland (2016-2017), and Labrador-Davis Strait (2003-2018) found no matches with individuals photographed in this study.

## Conclusions

Many interactions between whales and fisheries are detrimental to the individuals involved, as whales can face consequences including by-catch, entanglement and retaliation by fishers (3). Whales that change their foraging behavior to associate with trawlers may also become dependent and vulnerable to changes in fishing effort or practices. Although some studies have shown that associations with fisheries have benefitted whales (e.g., killer whales in the Crozets; (5, 29)), the energetic implications and risks of interactions for whales described here are unknown and need further study.

Our observations and analysis suggest that sperm whales in the study area have continued and possibly increased their associations with trawl fisheries in recent decades and modified their foraging behavior. Meanwhile northern bottlenose whales, not previously documented to interact with trawlers in the study area, have learnt to take advantage of novel feeding opportunities, as they have in other regions. Given the evidence that social learning can increase the incidence of whale depredation and foraging associations with fisheries (25), further study is needed to understand whether and how social transmission may be a factor in this area.

Data for this study were collected by the first author while working as a NAFO fisheries observer, demonstrating that fisheries observers can effectively record additional data on whale interactions which can be valuable for research and management. Information on fishery interactions with whales is otherwise rarely available and improving observer reporting requirements for international fisheries could provide critical data necessary for supporting ecosystem-based fisheries management.

## Author Contributions

UO and JG contributed to conception and design of the study. UO wrote the first draft of the manuscript. LF contributed to the statistical analysis, writing and figures. All authors contributed to manuscript revision, read, and approved the submitted version.

## Acknowledgements

We would like to thank the Playa Menduiña Dos’ crew and I. Marigómez for their support in this project. UO began her career as a fisheries observer on the Villa de Pitanxo, where she first saw whales following a vessel, and initiated the idea for this study. Sadly, the Villa de Pitanxo sank February 15, 2022, fishing in the same waters where this study took place, killing twenty-one, including the observer, and leaving three survivors. We would like to acknowledge the contributions of fishers and observers everywhere to increasing our knowledge and understanding of rare marine species and ecosystems.

## Supplementary Data

