## Supplemental Tables and Figures for "Sperm and northern bottlenose whale interactions with deep-water trawlers in the western North Atlantic"

**Supplementary Data**

### Table S1. Summary of sperm whale photo id data, showing date, haul number, time, fishing area, trawler activity and speed for each encounter. Trawler activity codes: (H) Hauling (P/S) Preparing/Shooting (T) Towing

| **Name** | **ID features** | **Date** | **Haul #** | **Time** | **Fishing Area** | **Activity** | **Associates** | **Distance between encounters (km)** |
| --- | --- | --- | --- | --- | --- | --- | --- | --- |
| **SPARROW** | scar behind dorsal fin, two little scars close to the blow hole and fluke | 7/8/2007 | 44 | 17:00 | Playa Nueva | H | BREIXO, SCRATCHY | **-** |
|  |  | 10/8/2007 | 48 | 10:45 | Playa Nueva | H | NEBOA, BREIXO | **5** |
|  |  | 14/08/2007 | 55 | 9:00 | Playa Nueva | H | SCRATCHY | **9** |
|  |  | 15/08/2007 | 58 | 19:30 | Playa Nueva | H |  | **58** |
|  |  | 11/9/2007 | 125 | 18:00 | Playa Nueva | P/S |  | **16** |
| **SCRATCHY** | scratch in left lateral | 7/8/2007 | 44 | 17:00 | Playa Nueva | H | SPARROW, BREIXO | **-** |
|  |  | 14/08/2007 | 55 | 9:00 | Playa Nueva | H | SPARROW | **4** |
| **BREIXO** | right lateral scar, snout scar and fluke | 7/8/2007 | 44 | 17:00 | Playa Nueva | H | SPARROW, SCRATCHY | **-** |
|  |  | 10/8/2007 | 48 | 10:45 | Playa Nueva | H | NEBOA, SPARROW | **5** |
|  |  | 14/08/2007 | 56 | 9:45 | Playa Nueva | P/S |  | **7** |
|  |  | 14/08/2007 | 56 | 11:00 | Playa Nueva | T | NEBOA | **13** |
|  |  | 27/08/2007 | 86 | 10:15 | Carson Canyon | P/S | FANECA, IBO | **235** |
| **NEBOA** | scars in both laterals and fluke | 8/8/2007 | 45 | 18:15 | Playa Nueva | H |  | **-** |
|  |  | 8/8/2007 | 46 | 20:15 | Playa Nueva | T |  | **8** |
|  |  | 10/8/2007 | 48 | 10:45 | Playa Nueva | H | BREIXO, SPARROW | **113** |
|  |  | 10/8/2007 | 49 | 22:00 | Playa Nueva | H |  | **1** |
|  |  | 13/08/2007 | 54 | 18:15 | Playa Nueva | T |  | **57** |
|  |  | 14/08/2007 | 56 | 11:00 | Playa Nueva | T | BREIXO | **50** |
|  |  | 14/08/2007 | 56 | 18:15 | Playa Nueva | T |  | **36** |
|  |  | 14/08/2007 | 56 | 21:00 | Playa Nueva | H |  | **12** |
| **FANECA** | left lateral scar, fluke | 27/08/2007 | 86 | 10:15 | Carson Canyon | P/S | BREIXO, IBO | **-** |
|  |  | 28/08/2007 | 88 | 21:05 | Carson Canyon | H |  | **59** |
|  |  | 29/08/2007 | 90 | 9:30 | Carson Canyon | T |  | **54** |
|  |  | 29/08/2007 | 90 | 14:45 | Carson Canyon | T |  | **31** |
|  |  | 29/08/2007 | 91 | 21:15 | Carson Canyon | T |  | **5** |
| **IBO** | scar in dorsal fin and three little depressions in the surface of the head | 27/08/2007 | 86 | 10:15 | Carson Canyon | P/S | FANECA, BREIXO | **-** |
|  |  | 9/9/2007 | 121 | 16:30 | Playa Nueva | H |  | **276** |
| **MARMU** | fluke | 8/8/2007 | 45 | 13:23 | Playa Nueva | T |  | **-** |
| **TOR** | scar in blow hole | 29/07/2007 | 25 | 11:10 | Playa Nueva | H |  | **-** |
| **NAT** | fluke | 28/07/2007 | 23 | 10:44 | Playa Nueva | H |  | **-** |
| **NOAH** | scar in left lateral and fluke | 29/08/2007 | 91 | 21:15 | Carson Canyon | T | FANECA | **-** |

### Figure S1. Associations between identified sperm whales. The arrows show the presence of both individuals together near the vessel, and the number of times the whales were seen together. *Breixo* (middle) was seen with all the other animals.

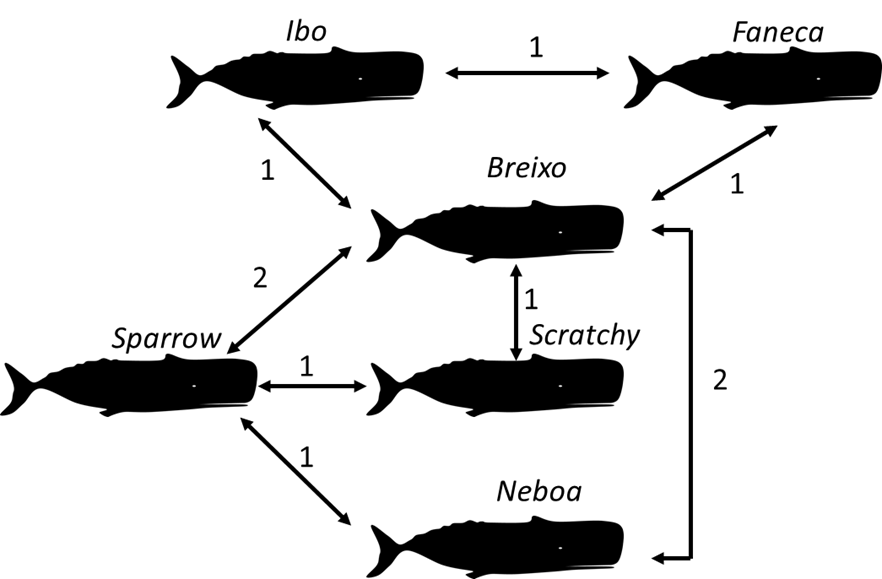

### Figure S2. Location of sightings of individual sperm whales in (A) the entire study area and (B) an inset of concentrated sightings in the Flemish Pass.

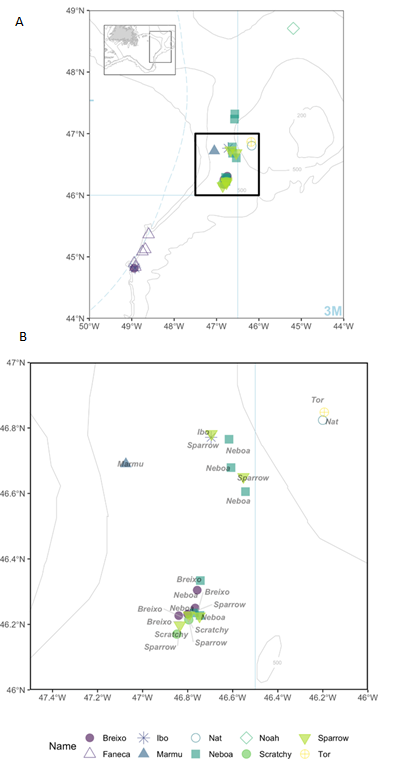
